# Introduction, spread, and evolutionary dynamics of non-native spiders in North America

**DOI:** 10.64898/2025.12.16.694649

**Authors:** Chao Tong, Miao Li, Zhisheng Zhang

**Affiliations:** School of Life Sciences, Arizona State University, Tempe, AZ 85281, USA; Center for Bioinformatics and Computational Biology, University of Delaware, Newark, DE 19713, USA; School of Life Sciences, Southwest University, Chongqing 400715, China

**Keywords:** Global introduction, Non-native species, North America, Spiders

## Abstract

Tracing the introduction, spread, and evolution of non-native species is crucial for risk assessment and management. Spiders represent one of the largest groups of terrestrial arthropod predators, and an increasing number of species have been introduced globally, posing growing challenges to local biodiversity and ecosystem stability. Yet, their global introduction routes, spatial diversity patterns, and evolutionary dynamics remain poorly understood. Here, we conducted a global-scale survey of 53,172 taxon records and updated the checklist of North American spiders to 5,401 species, including 118 non-native species. By integrating historical records with species occurrence data, we reconstructed the geographic origins of non-native species and found that most originated from Asia and Europe, consistent with long-standing transcontinental trade and intensive human activities. Mapping the contemporary distributions of all non-native species revealed at least six richness hotspots, such as Los Angeles and San Francisco metropolitan areas, underscoring strong associations between non-native spider accumulation and population density, trade intensity, and human-mediated dispersal pathways. By analyzing genomes of 45 native and five non-native spider species within North America, we detected a set of metabolic, stress tolerance and immune defense genes under positive selection and rapid evolution in non-native species, suggesting that putative functional innovations may contribute to their establishment, expansion, and ongoing local adaptation. Altogether, our study provides the most comprehensive survey to date of introduction routes, spatial dynamics, and evolutionary signatures of non-native spiders in North America, and highlights putative mechanisms that may facilitate arthropod introductions under global change.

## Introduction

While the dispersal of species beyond their native geographic ranges is a natural biogeographic process, human-mediated globalization has amplified its frequency and scale to unprecedented levels, dramatically reshaping global biodiversity (1, 2). Most non-native species are benign (3). However, a subset becomes invasive, posing substantial threats to local ecosystems and agriculture. Reconstructing the introduction routes of non-native species is fundamental for risk assessment and management, yet this task is often hindered by sparse, geographically biased, or entirely absent historical records, especially for hyperdiverse invertebrate groups (4–6).

Spiders (Araneae), a globally distributed and ecologically successful group of terrestrial arthropod predators, are frequently intercepted in global trade (4, 6–8), but a systematic, continental-scale assessment of their non-native fauna has never been conducted. To understand the ecological or evolutionary consequences of introductions, a systematic inventory of non-native species and their geographic origins is a foundational imperative.

Following establishment in a new continent, non-native species populations are subject to novel selective pressures, from different climates, competitors, and prey, which can lead to rapid local adaptation. Advances in comparative genomics provide a robust framework to identify signatures of natural selection and provide insights into evolutionary dynamics associated with post-introduction or invasion (9, 10). North America represents a major hotspot of biological introductions, yet past research has focused largely on vertebrates and several arthropod groups (6). Consequently, the geographic origins, introduction routes, spread and genomic evolutionary dynamics of non-native spiders remain largely unknown. In this study, we aim to address three questions: (i) Which spider species in North America are non-native? (ii) Where did these non-native species come from? (iii) How do they evolve following introduction? We conducted a large-scale survey by analyzing more than 50,000 taxon records from the World Spider Catalog (WSC) and occurrence records from the Global Biodiversity Information Facility (GBIF). We further performed comparative genomic analyses of native species and non-native species collected within North America and identify genomic evolutionary dynamics associated with local adaptation following introduction.

## Results

We examined 53,172 taxon records from the WSC and GBIF databases and determined 5,401 spider species occurring in North America (Fig. 1A; Dataset S1), comprising 5,283 native and 118 non-native species across 76 families. We found that non-native species were disproportionately concentrated in Theridiidae (N = 16), Salticidae (N = 15), Linyphiidae (N = 11), Gnaphosidae (N = 10), Oonopidae (N = 9) and Pholcidae (N = 7), which together accounted for the majority of introductions. By analyzing the non-native species at the global scale, we also found the top concentrated families of non-native spiders were from Theridiidae (N = 52), Salticidae (N = 50), Linyphiidae (N = 30) (Dataset S2). We further quantified the proportion of non-native species within each family and found several previously unrecognized non-native lineages in North America, including Cithaeronidae, Desidae, and Scytodidae (Fig. 1A, Dataset S3).

**Figure 1.**
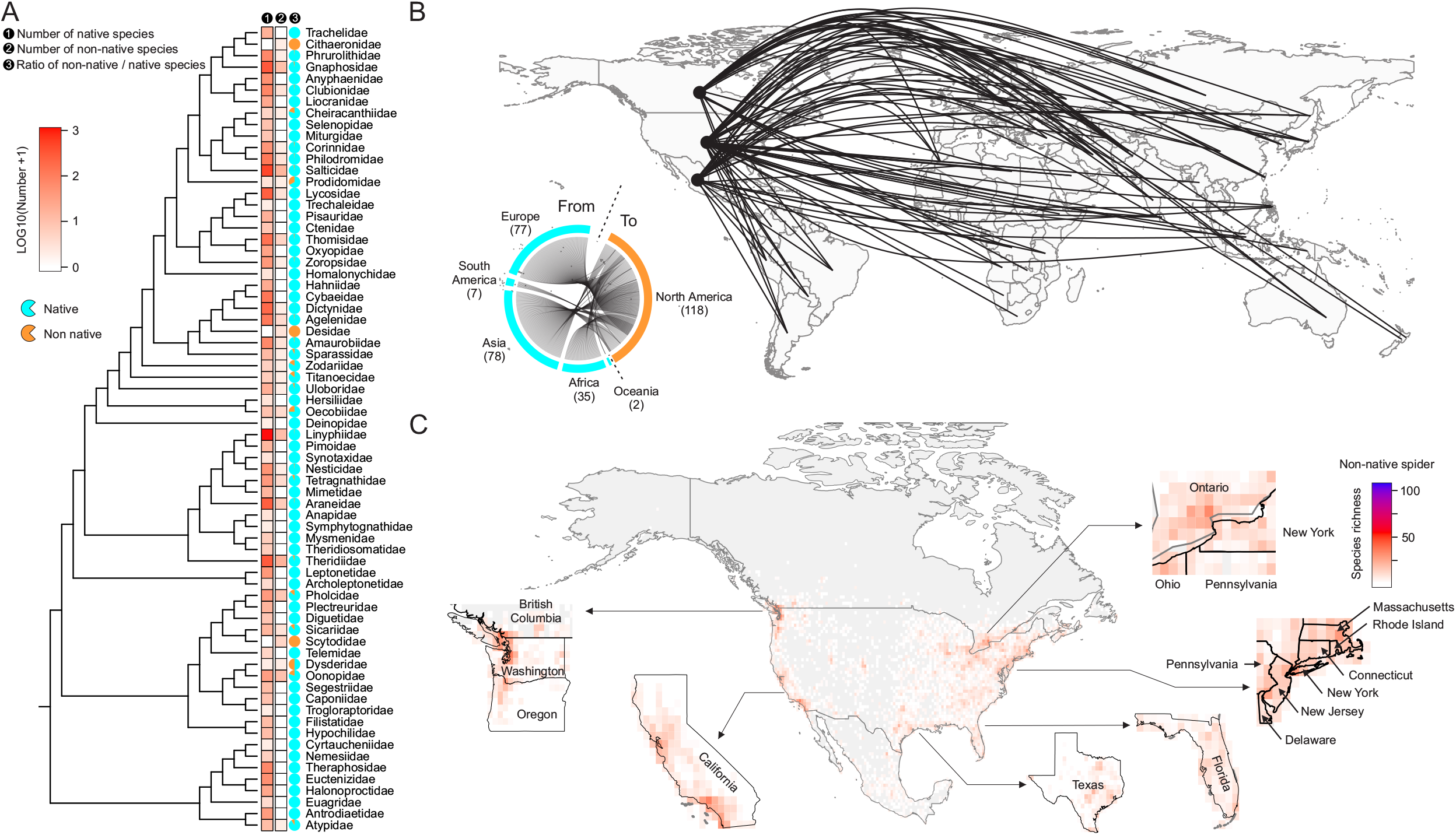
Non-native spiders in North America. (A) Family-level phylogeny is trimmed from a published phylogeny (29). Heatmaps depicting the number of native and non-native species in each family. Pie diagrams depicting the ratio of non-native / native species in each family. (B) A map showing the origins of non-native spider species introduced to North America. Each line represents the inferred origin of a single species. Circular plot depicting the number of non-native spider species from five different continents. (C) Spatial pattern of non-native species richness in North America. Several hotspots of non-native species richness were extracted from the map and highlighted.

We further assembled historical records and traced each non-native species’ geographic origin. We found that most non-native spiders in North America originated from Asia (n = 78) and Europe (n = 77) (Fig. 1B). We collected the contemporary distribution data of all 118 non-native species and identified at least six hotspots having exceptional non-native species richness.

These included the San Francisco Bay area and Los Angles metropolitan areas in California, the Vancouver - Seattle of the Pacific Northwest, the New England region and Toronto metropolitan areas in Northeastern America, as well as eastern Texas and peninsular Florida, each exhibiting remarkedly elevated accumulation of established non-native lineages (Fig. 1D).

We finally analyzed the publicly available genomic datasets of 45 native and five non-native spider species within North America (Dataset S4). We detected a set of genes under positive selection with significance in non-native species, such as 36 genes in *Parasteatoda tepidariorum* (Fig. 2A, Dataset S5). Among these positively selected genes (PSGs), we identified several PSGs shared by multiple non-native spiders, including an ion transport gene, *Chloride Channel Accessory 4* (*CLCA4*) in *Pholcus phalangioides* and *Metaltella simoni*, a spermatogenesis gene, *Paramyosin* (*Prm*) in *Metaltella simoni* and *Parasteatoda tepidariorum*, a stress-responsive transcription factor gene, *cAMP-responsive element-binding protein 3* (*CREB3*) in *Pholcus phalangioides* and *Falconina gracilis* (Fig. 2B). By comparing the molecular evolutionary rates of genes between native and non-native spider species, we identified 148 genes with accelerated patterns of molecular evolution in non-native spiders (rapidly evolving genes, REGs) and 106 genes with decelerated patterns of molecular evolution in non-native spiders (slowly evolving genes, SEGs) (Fig. 2C, Dataset S6). These REGs are associated with several key functions, including *Probable chitinase 10* (*Chit10*) involved in digestion and immune defense, *DNA mismatch repair protein Msh2* (*MSH2*) involved in maintaining genome integrity, *Prm* and *chorion peroxidase* (*Pxt*) associated with reproduction, *heat shock protein 75 kDa* (*TRAP1*) and *superoxide dismutase [Mn]* (*SOD2*) involved in thermal and oxidative tolerance, *ATP-binding cassette sub-family D member 2* (*ABCD2*) associated with lipid metabolism and detoxification, *cytochrome P450 2J4* (*CYP2J4*) involved in xenobiotic metabolism.

**Figure 2.**
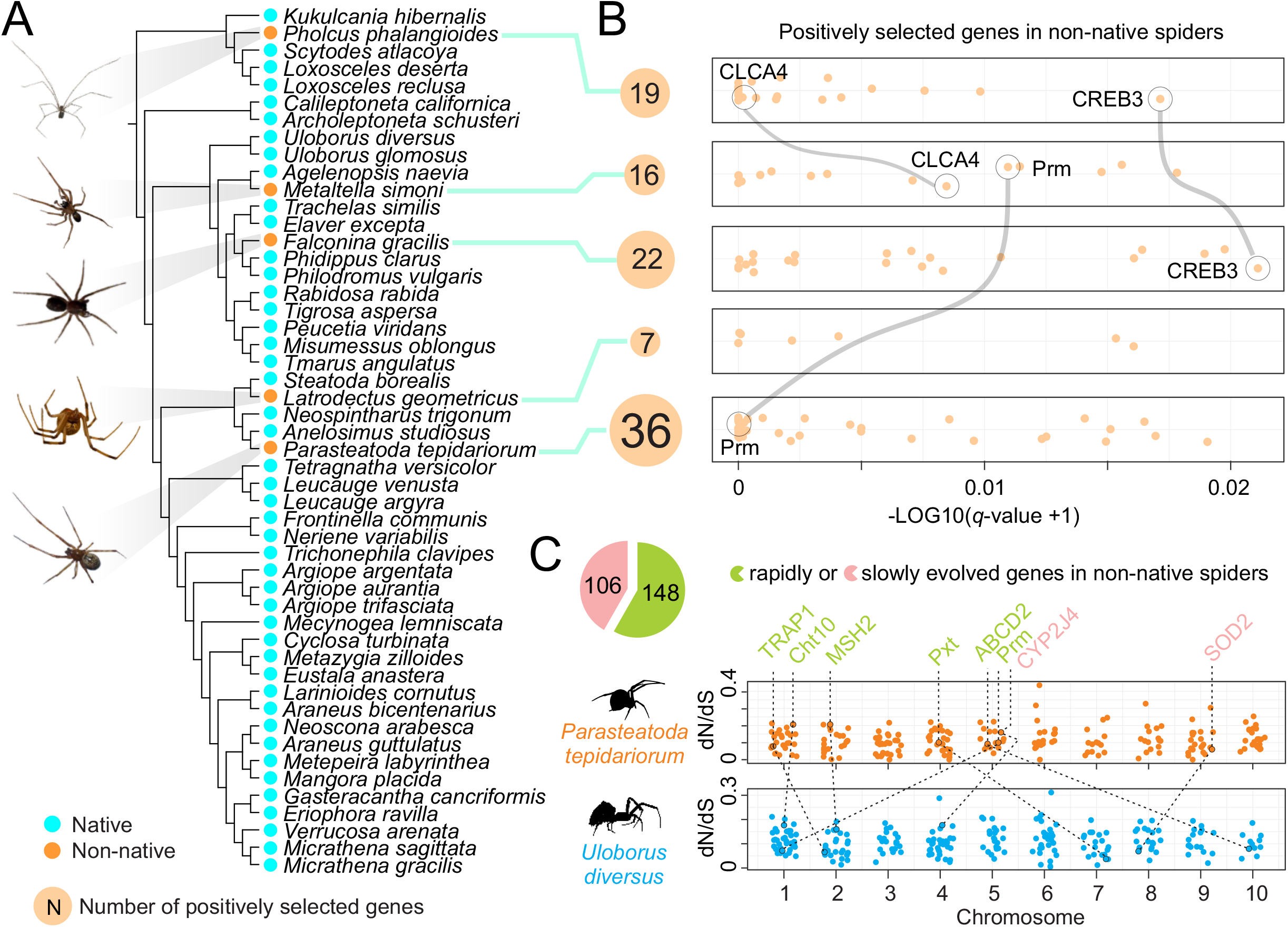
Evolutionary dynamics of non-native spiders. (A) Genome-scale phylogeny of 45 native and five spider species within North America. Cyan dots represent native species and light orange dot represent non-native species. Photos of five non-native species were extracted from iNaturalist (https://www.inaturalist.org/). (B) Sand-colored bubbles depicting the number of positively selected genes in each non-native spider species. Dot plots depicting the positively selected genes, and x-axis represents the value of *q* values estimated by aBS-REL after log transformation. Shared positively selected genes were highlighted. (C) Pie diagram depicting the number of genes under acceleration (yellow-green) or deceleration (pink) in non-native species relative to native species. Dot plots depicting the distributions of accelerated or decelerated genes on each chromosome of representative native species (*Parasteatoda tepidariorum*) and non-native species (*Uloborus diversus*).

## Discussion

How many spider species are in North America? Previous studies had suggested that approximately 3700 species were documented in this region (11, 12). However, species diversity and richness have continued to increase, while the checklist has remained largely outdated. Our study refines estimates of species diversity and taxonomic composition to date and updated the checklist of North American spiders to 5,401 species within 76 families. We further determined 118 non-native species, which have become established across the continent. How and where did they come to North America? By integrating our findings with historical evidence suggests that early introductions of non-native spiders were closely associated with transatlantic trade, transportation, and human migration between Europe and North America, during which spiders were inadvertently transported with cargo, timber, and ornamental plants (5). In more recent decades, the intensification of global trade and increasing exchanges between North America and Asian regions likely facilitated additional introductions (13), this pattern is consistent with our reconstruction of introduction routes showing Asia and Europe as primary origins of non-native spiders. This finding also aligned with evidence from invasive insects studies, indicating that Europe represents a dominant source region for non-native arthropods in North America (14). Analyses of contemporary distribution and richness patterns reveal that non-native spiders are disproportionately concentrated in major metropolitan areas and along both the eastern and western coasts, where international trade and human activity are most intense. This spatial pattern is consistent with localized establishment and potential local adaptation, rather than widespread continental expansion.

Nonetheless, increasing richness in parts of central North America suggests that ongoing intensification of human activities may promote future range expansion.

The rapid advances in comparative genomics allow the investigation of evolutionary dynamics associated with ongoing local adaptation in non-native spiders. Intriguingly, although *Latrodectus geometricus* is currently recognized as an invasive species with notable ecological impact (15), we detected relatively few PSGs. In contrast, *Parasteatoda tepidariorum* exhibited a relatively larger number of PSGs, which may be related to differences in ecological tolerance, as *L. geometricus* is generally restricted to warmer environments, whereas *P. tepidariorum* has a much broader distribution and can tolerate a wide range of thermal conditions. We further compared rates of molecular evolution of genes between native and non-native spiders and found that many genes associated with energy metabolism evolved more rapidly in non-native spiders, potentially reflecting strong environmental adaptability. In addition, we identified rapidly evolving genes related to detoxification, reproduction, immune functions and stress tolerance in non-native spiders, which may indicate a genetic basis for exploiting diverse prey resources.

Collectively, these results reveal the introduction routes, spatial patterns, and putative evolutionary dynamics of non-native spiders in North America and provide insights into invasion risk assessment and biodiversity conservation management.

## Materials and Methods

### Taxon and occurrence record retrieval

We retrieved the distribution datasets and relevant literatures of all described spider species from the World Spider Catalog database (WSC, https://wsc.nmbe.ch/) as of 25 December, 2024. In addition, we conducted thorough searches for occurrence records of described spider species in the Global Biodiversity Information Facility (GBIF, https://www.gbif.org/). We documented taxonomy, geographic distribution and literature that corresponded to each spider species. Finally, we assembled a curated list comprising non-native spider species with records of description “introduced” or “invasive” as the search terms in above documented datasets.

### Estimating geographic origins of non-native species

We manually checked all the relevant literatures of candidate non-native species and the historic records of spatial distributions in WSC database, determined the geographic origins and recent introduced ranges. For example, past literatures and records indicated that Pholcus manueli is originated from Kazakhstan, Turkmenistan, Russia (Far East), China, Korea, Japan, and introduced to USA.

### Spatial distribution of non-native species richness

We downloaded the occurrence records of non-native spider species from GBIF database and retained the records specific to North America mainland. All the distribution records were gridded into cells of 1 x 1 degree size (∼111 x 111 km^2^) on the map.

### Genome data collection

We explored the whole genome data of spider from online databases, such as NCBI Genome (https://www.ncbi.nlm.nih.gov/genome) and GigaDB (http://gigadb.org/), as of 25 December, 2024. In addition, we manually checked the sample detail information of each published spider genome and retained the spider genomes based on the specimens from North America. Finally, we employed the BUSCO pipeline (16) to assess the completeness of genome contents of these spider species based on the arachnida_odb10 single-copy orthologous gene set from OrthoDB v12 (https://www.orthodb.org), which comprised 2,934 conserved genes.

### Transcriptome data collection and assembly

We extended the available genomic dataset by including transcriptomes of spiders collected from North America. We downloaded the raw transcriptome sequencing reads of North America spiders from NCBI SRA database (https://www.ncbi.nlm.nih.gov/sra), and performed *de novo* transcriptome assembly using rnaSPAdes (17). Further, we used TransDecoder (https://github.com/TransDecoder/TransDecoder) to predict gene models in each assembled transcriptome. Finally, we also used BUSCO pipeline to assess the quality of transcriptome assemblies.

### Phylogenetic tree reconstruction

We included a genome of *Centruroides vittatus* (GCF_030686945.1) for phylogenetic tree reconstruction as the requirement of an outgroup species. We assembled a dataset including previously identified BUSCO gene repertoire within two categories “Complete and single-copy BUSCO” and “Complete and duplicated BUSCO” of 50 spider species. We further inferred one-to-one single-copy genes across all 50 species using OrthoFinder (18).

We used a phylogenomic approach to reconstruct the phylogeny of 50 spider species, along with an outgroup species, based on a dataset of amino acid (AA) sequences corresponding to a pooled set of 1:1 single-copy orthologs. Further, we performed AA sequence alignment using MAFFT (19), removed gaps using trimAL (20), and assembled a concatenated dataset that included all 1:1 single-copy orthologs with a minimum length of 200 AA. Finally, we used ModelFinder (21) to determine best-fit model of sequence evolution and constructed the maximum likelihood (ML) phylogenetic tree using IQ-TREE2 (22) with 1000 bootstrap replicates.

### Gene Orthology

We extended and explored the orthologous relationship of protein-coding genes across 50 spider species using FastOMA (23), because BUSCO gene repertoire only included curated single-copy orthologs from OrthoDB dataset. Briefly, we compiled protein-coding gene dataset for each species and amalgamated them into a local pooled protein database. We performed parallel search using DIAMOND (https://github.com/bbuchfink/diamond) and identified putative orthologous groups (OGs). For each 1:1 ortholog pair, we selected the longest gene associated with curated OG as putative ortholog for each species.

### Analysis of positive selection in non-native spiders

To determine whether natural selection (i.e. positive selection) associated with post-introduction, we sought to detect the signal of positive selection in shared orthologs of 50 species across the phylogeny. Specifically, we prepared the codon alignments of shared orthologs by 50 species, which derived from amino acid alignments and corresponding DNA sequences using PAL2NAL v.14 (-no gap) (24). We retained the codon alignments with a minimum length of 50 codons, and prepared the corresponding tree for each ortholog by pruning the genome-scale phylogeny using R package, phytools (25). We applied an adaptive Branch-Site Random Effects Likelihood model (aBS-REL) (26) in HyPHY (27). For each shared ortholog, aBS-REL was used to test whether the terminal branch corresponding to the focal species experienced episodic positive selection, allowing the nonsynonymous-to-synonymous substitution ratio (dN/dS, ω) to vary both across sites and across branches. In addition, likelihood ratio test (LRT) was performed for all branches in the gene tree, and p-values were corrected for multiple testing across branches using the Holm–Bonferroni correction in aBS-REL. Genes were classified as being under positive selection in the focal species if q-value for the corresponding branch was less than 0.05, also known as positively selected genes.

### Analysis of shifts in evolutionary rate between non-native and native spiders

To characterize signatures of selection pressure in native and non-native spider species, we applied a model that constrains the ratio of non-synonymous to synonymous rate (dN/dS, ω) in two groups and then employs a likelihood ratio test. Specifically, ω rates were estimated for two discrete categories including non-native and native species under the the Muse-Gaut (MG94) rate matrices (28) in HyPHY (27), and then compared nested models with constrained relationships among them. This approach allowed to detect the shift of selective pressure associated with post-introduction (alternative model, H1; one ω rate represents native spiders: ω_native_; another ω rate represents non-native spiders: ω_non-native_) and compared to the null model (H0) that assumed all branches have the same ω rate. We used FitMG95 (https://github.com/veg/hyphy-analyses/tree/master/FitMG94) in HyPHY to repeat the rest ten times for each gene and results in ten likelihood scores for each H0 and H1. We then constructed the log-likelihood ratio score for each gene (ΔlnL) as: ΔlnL = 2(lnL H1-lnL H0) = 2(Maxi-10 (LnL H1) - Maxi-10 (LnL H0)) and employed the LRT. Further, we corrected the reported p-values for multiple comparisons by computing q-value and considered these genes experienced shift in selective pressure. Finally, we defined these significant genes (q-value < 0.05) with high ω rate in non-native spiders than native spiders as rapidly-evolving genes, and otherwise as slowly evolving genes.

## Data, Materials, and Software Availability

All code and scripts are available on Github (https://github.com/jiyideanjiao/spider_introduction_north_america). All study materials, detail methods are described in Supporting Information.

## Reference

1. C. S. Elton, The ecology of invasions by animals and plants, 2nd Ed. (Springer Nature, 2020).

2. H. Seebens, et al., No saturation in the accumulation of alien species worldwide. Nat. Commun. 8, 14435 (2017).

3. T. M. Blackburn, et al., A unified classification of alien species based on the magnitude of their environmental impacts. PLoS Biol. 12, e1001850 (2014).

4. A. Chuang, J. F. Deitsch, D. R. Nelsen, M. I. Sitvarin, D. R. Coyle, The Jorō spider (Trichonephila clavata) in the southeastern U.S.:an opportunity for research and a call for reasonable journalism. Biol. Invasions 25, 17–26 (2023).

5. P. E. Hulme, Unwelcome exchange: International trade as a direct and indirect driver of biological invasions worldwide. One Earth 4, 666–679 (2021).

6. B. M. Marshall, et al., Mapping the global dimensions of US wildlife imports. Curr. Biol. 35, 3959–3972.e4 (2025).

7. B. M. Marshall, et al., Searching the web builds fuller picture of arachnid trade. Commun. Biol. 5, 448 (2022).

8. H. Seebens, et al., Biological invasions: a global assessment of geographic distributions, long-term trends, and data gaps. Biol. Rev. Camb. Philos. Soc. 100, 2542–2583 (2025).

9. J. Hudson, S. D. Bourne, H. Seebens, M. A. Chapman, M. Rius, The reconstruction of invasion histories with genomic data in light of differing levels of anthropogenic transport. Philos. Trans. R. Soc. Lond. B Biol. Sci. 377, 20210023 (2022).

10. D. B. Stern, C. E. Lee, Evolutionary origins of genomic adaptations in an invasive copepod. Nat. Ecol. Evol. 4, 1084–1094 (2020).

11. R. A. Bradley, Common Spiders of North America. (2012).

12. P. Paquin, D. J. Buckle, N. Dupérré, C. D. Dondale, Checklist of the spiders (Araneae) of Canada and Alaska. Zootaxa 2461, 1 (2010).

13. D. S. Jacks, J. P. Tang, Trade and immigration, 1870-2010. (2018).

14. C. Bertelsmeier, et al., Temporal dynamics and global flows of insect invasions in an era of globalization. Nat. Rev. Biodivers. 1, 90–103 (2025).

15. A. C. Wahlberg, R. Antoniazzi, C. M. Schalk, Patterns of the introduction, spread, and impact of the brown widow spider, Latrodectus geometricus (Araneae: Theridiidae), in the Americas. J. Arachnol. 51 (2023).

16. F. A. Simão, R. M. Waterhouse, P. Ioannidis, E. V. Kriventseva, E. M. Zdobnov, BUSCO: assessing genome assembly and annotation completeness with single-copy orthologs. Bioinformatics 31, 3210–3212 (2015).

17. E. Bushmanova, D. Antipov, A. Lapidus, A. Prjibelski, rnaSPAdes: a de novo transcriptome assembler and its application to RNA-Seq data. GigaScience 8 (2018).

18. D. M. Emms, S. Kelly, OrthoFinder: phylogenetic orthology inference for comparative genomics. Genome Biol. 20, 238 (2019).

19. K. Katoh, D. M. Standley, MAFFT multiple sequence alignment software version 7: improvements in performance and usability. Mol. Biol. Evol. 30, 772–780 (2013).

20. S. Capella-Gutiérrez, J.M. Silla-Martínez, T. Gabaldón, trimAl: a tool for automated alignment trimming in large-scale phylogenetic analyses. Bioinformatics 25, 1972–1973 (2009).

21. S. Kalyaanamoorthy, B. Q. Minh, T. K. F. Wong, A. von Haeseler, L. S. Jermiin, ModelFinder: fast model selection for accurate phylogenetic estimates. Nat. Methods 14, 587–589 (2017).

22. B. Q. Minh, et al., IQ-TREE 2: New Models and Efficient Methods for Phylogenetic Inference in the Genomic Era. Mol. Biol. Evol. 37, 1530–1534 (2020).

23. S. Majidian, et al., Orthology inference at scale with FastOMA. Nat. Methods 1–4 (2025).

24. M. Suyama, D. Torrents, P. Bork, PAL2NAL: robust conversion of protein sequence alignments into the corresponding codon alignments. Nucleic Acids Res. 34, W609–12 (2006).

25. L. J. Revell, phytools 2.0: an updated R ecosystem for phylogenetic comparative methods (and other things). PeerJ 12, e16505 (2024).

26. M. D. Smith, et al., Less is more: an adaptive branch-site random effects model for efficient detection of episodic diversifying selection. Mol. Biol. Evol. 32, 1342–1353 (2015).

27. S. L. Kosakovsky Pond, et al., HyPhy 2.5—A Customizable Platform for Evolutionary Hypothesis Testing Using Phylogenies. Mol. Biol. Evol. 37, 295–299 (2020).

28. S. V. Muse, B. S. Gaut, A likelihood approach for comparing synonymous and nonsynonymous nucleotide substitution rates, with application to the chloroplast genome. Mol. Biol. Evol. 11, 715–724 (1994).

29. J. O. Wolff, et al., Stabilized morphological evolution of spiders despite mosaic changes in foraging ecology. Syst. Biol. 71, 1487–1503 (2022).

